# Effective Variations of related Physicochemical Properties of Nucleotides Leading to Amino Acids for Characterizing Genes and Proteins

**DOI:** 10.1101/332171

**Authors:** Antara Sengupta, Pabitra Pal Choudhury

## Abstract

The aim of this paper is to make quantitative analysis of the properties which is really being carried from DNA sequence and finally landing up to the properties of a protein structure through its primary protein sequence. Thus, the paper has a theory which is applicable for any arbitrary DNA sequence whether it is of various species or mutated data or a bunch of genes responsible for a function to be occurred. Irrespective to genes of any families, species, wild type or mutated, our paper here gives a standard model which defines a mapping between physicochemical properties of any arbitrary DNA sequence and physicochemical properties of its amino acid sequence. Experiments have been carried out with PPCA protein family and its four homologs PPC(B E) which establishes that DNA sequence keeps its signature even after its translation into the corresponding amino acid sequence.

## Introduction

According to medical definition of Central Dogma – “Genetic information is coded in self replicating DNA and undergoes unidirectional transfer to messenger RNAs in transcription which acts as template for protein synthesis in translation”[22]. The definition itself implies that the canonical genetic code maps DNA codons onto corresponding amino acids and that mapping follow predefined protocols. There are 61 normal codons and 3 stop codons. Each codon codes for a specific Amino acid and each amino acid is responsible for a specific protein synthesis. As a result, it is clear that they do not have one to one mapping between them, instead some amino acids are mapped with more than one codon and degeneracy takes its birth which initiates ‘multiplet structure’ of genetic code [6]. As more than one codon can code for a specific amino acid so there must be specific protocols that are being adopted by nature for selection of codons to code different amino acids in a primary protein sequence. Several research papers are reported where researchers demand existence of relationships between the codons and the physical chemical (physicochemical) properties of the coded amino acids. Our literature survey observes that although the degeneracy is found primarily at the third position of a codon, the nucleotides at the first two positions are mostly sufficient to specify a unique amino acid. According to Lagerkvist[1][2] ‘‘codons may be read according to the two out of three principle that relies mainly on the Watson–Crick base pairs formed with the first two codon positions, while the miss paired nucleotides in the third codon and anticodon wobble positions make a comparatively smaller contribution to the total stability of the reading interaction’’.

In this paper we have tried to investigate the physicochemical properties of DNA sequence that keep their signatures embedded while translated into the protein synthesis. Thus, here we have clearly defined a mapping between physicochemical properties of any arbitrary DNA sequence and physicochemical properties of its amino acid sequence. Each DNA sequence in our universe starts its journey with some specific physicochemical properties. Pyrimidine (Y=C/T) has one heterocyclic ring, whereas, Purine (R=A/G) has two heterocyclic rings, while in RNA sequence T becomes U. As the structure of a DNA is a double helix whose backbone is composed of sugars bound to phosphates, the bases face inward with a purine always binding to a pyrimidine. In this way, the helix bases are complementarily paired through hydrogen bonds. A makes bond with U and C bonds with G. U or A have 2 hydrogen bonds and they are weak(W), whereas, C or G have 3 hydrogen bonds and are strong(S). keto (K = U or G) or amino (M = A or C) have 6 keto and 6 amino bases respectively. If we take two same bases at a time out of A, C, G, and T, we have four combinations: AA, CC, GG and TT. The structures have been shown in the figure 1. Consequently, we obtain the Double Nucleotide or Dual Nucleotide set (DN set)**[19]** of 16 dual nucleotides: {AG,GA,CU,UC,AC,CA,GU, UG,AU,UA,CG,GC,AA,CC,GG,UU} which are responsible to code first two bases of 64 codons of Genetic Code table by introducing another base at third position in it.

**Fig 1:**
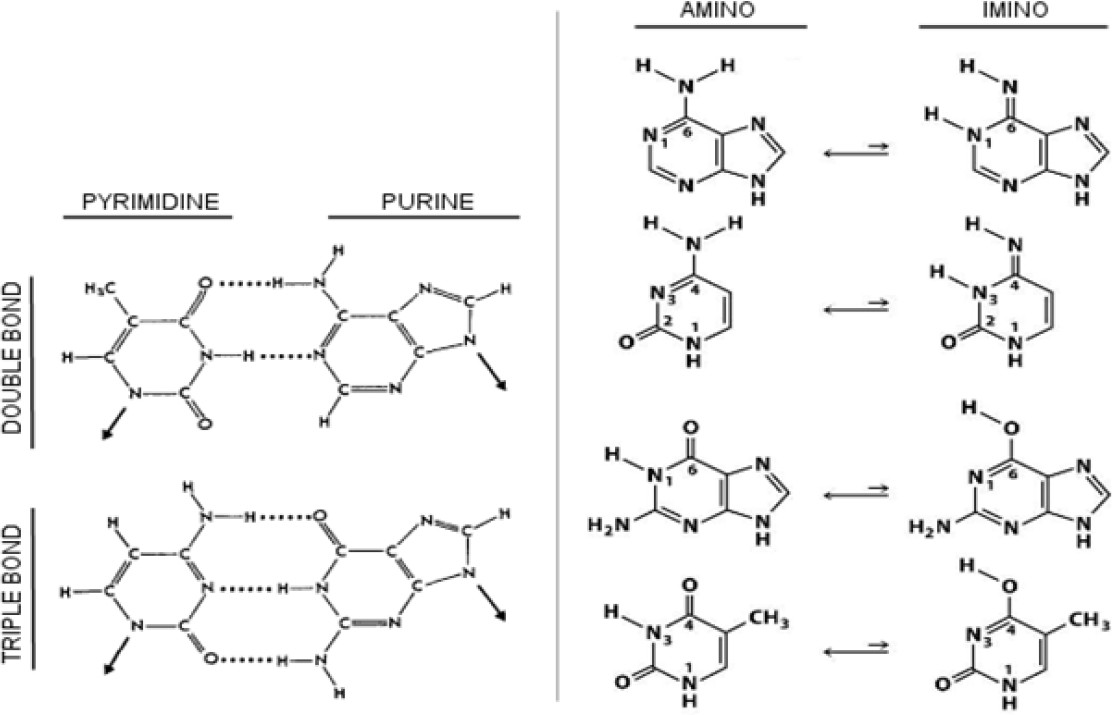
Structures of Amino/Keto, and Purine/Pyrimidine[15].

In this paper, the section of materials and methods has several sub sections. In first sub section the whole methodology has been described in detail. A standard theory has been tried to establish regarding the interrelationship between the physicochemical properties of DNA sequence and the physicochemical properties they achieve after translation. The second section contains the Applications and their results. Here in this section methodology has been applied to PPCA protein family and its four homologs and experiments have been carried out. Lastly, in the section 3, the whole facts and findings are concluded.

## 1. Materials and Methods

### 1.1 Binary interpretations of four bases of DNA sequence

> Mathematically DNA sequences (nucleotides) are symbolic sequences and may be interpreted as digital sequences to make computational analysis easier. As DNAs are one dimensional sequences, so they can be represented as binary strings. To uniquely identify 4 nucleotides, mathematically, log_2_ 4 = 2 bits (Binary Digits) are needed which are 00, 01, 10, 11. It is worth to state that there are total 24 numbers of possible ways we can assign U, A, C, G the values 00, 01, 10, 11. We know that A bonds with U and C bonds with G. This is analogous to lock and key arrangement meaning that one is complementary to the other. It is our aim to reflect this logic in binary representation too. Among those 24 possible combinations 8 possible combinations really obey this rule as shown below in the table 1. Here in this paper one such mapping is selected, T (U) =00, T (A) =11, T(C) =01, and T (G) =10. It is worth to state that as Watson Crick base pairs (guanine cytosine and adenine thymine) allow the DNA helix to maintain a stable regular helical structure, the binary representations have been chosen accordingly. The explanation has been given in the following paragraphs.

**Table 1:** Binary interpretations of four bases of DNA sequence.

| Binary Representation | U | A | C | G | Distance within Purine | Distance within Pyrimidine | Distance . In Keto | Distance In Amino | Distance In Strong H | Distance In Weak H |
| --- | --- | --- | --- | --- | --- | --- | --- | --- | --- | --- |
| Nemzer [3] | 0<br>0 | 1<br>0 | 0<br>1 | 1<br>1 | 1 | 1 | 1 | 2 | 1 | 1 |
| Our Manuscript | 0<br>0 | 1<br>1 | 0<br>1 | 1<br>0 | 1 | 1 | 1 | 1 | 2 | 2 |

For our analysis we concentrate on 3 pairs of chemical properties viz. 1. Purine (A,G) and Pyrimidine (C,T/U), 2. Strong H bond (C,G) and weak H bond (A,T/U), 3. Amino (A,C) and Keto (G,T/U). Group 1 and group 3 having distinct chemical properties. It is to be noted that within these groups hamming distances are consistently 1 for all 8 accepted possibilities of binary encodings shown in table**[2]**. Interestingly, group 2 consisting of strong H bond and weak H bond showing hamming distances of 2 consistently for all the 8 possibilities mentioned below in the table **2.** It is quite obvious to see that for providing the achieved storage in DNA (double helix bond) the distances are bigger, i.e. higher distance of 2 similar to the notion of opposite poles attract. Thus, our encodings (8 out of 24) carry more chemical/biological significances while the other sixteen encodings do not support these facts consistently.

**Table 2:** 8 possible 2 bit representations of 4 bases and their hamming distances

| Nucleotides | Distance between | 8 Possible 2 bit Representations of 4 bases and their hamming distances |  |  |  |  |  |  |  |  |  |  |  |  |  |  |  |
| --- | --- | --- | --- | --- | --- | --- | --- | --- | --- | --- | --- | --- | --- | --- | --- | --- | --- |
|  |  | VALUE | HD | VALUE | HD | VALUE | HD | ALU | HD | ALU | HD | ALU | HD | ALU | HD | ALU | HD |
| A | U and A | 00- | 2 | 00- | 2 | 11 | 2 | 11 | 2 | 01- | 2 | 01- | 2 | 10 | 2 | 10 | 2 |
| U | C and G | 11 | 2 | 11 | 2 | 00- | 2 | 00- | 2 | 10 | 2 | 10 | 2 | 01- | 2 | 01- | 2 |
| C | G and U | 01- | 1 | 10 | 1 | 01- | 1 | 10 | 1 | 00- | 1 | 11 | 1 | 00- | 1 | 11 | 1 |
| G | A and C | 10 | 1 | 01- | 1 | 10 | 1 | 01- | 1 | 11 | 1 | 00- | 1 | 11 | 1 | 00- | 1 |
|  | A and G |  | 1 |  | 1 |  | 1 |  | 1 |  | 1 |  | 1 |  | 1 |  | 1 |
|  | U and C |  | 1 |  | 1 |  | 1 |  | 1 |  | 1 |  | 1 |  | 1 |  | 1 |
|  | A and A |  | 0 |  | 0 |  | 0 |  | 0 |  | 0 |  | 0 |  | 0 |  | 0 |
|  | U and U |  | 0 |  | 0 |  | 0 |  | 0 |  | 0 |  | 0 |  | 0 |  | 0 |
|  | C and C |  | 0 |  | 0 |  | 0 |  | 0 |  | 0 |  | 0 |  | 0 |  | 0 |
|  | G and G |  | 0 |  | 0 |  | 0 |  | 0 |  | 0 |  | 0 |  | 0 |  | 0 |

Furthermore, although we have considered binary encoding U=00, A=11, C=01 and G=10, the hamming distances between all 4 bases will be same for all (8) possible encodings of nucleotide bases. The hamming distances are shown in the following table**[1].** Transitions make interchanges of either 2 rings for purine (A to G or G to A) or 1 ring for pyrimidine(C to U or U to C). So the hamming distance 1 specifies changes between bases of similar shapes. The encoding clearly distinguishes between pyrimidine (Y, U or C) and purine (R, A or G) bases. If we talk about inter instead of intra, viz. between U and A or C and G, they are called transversions and making the bonds. Thus, the Strong H and Weak H group of base pairs make changes between bases of different shapes or structures. Moreover Watson Crick base pairs (guanine cytosine and adenine thymine) allow the DNA helix to maintain a regular helical structure. The hamming distance 2 satisfies this fact in the sense of opposite poles attract as described earlier. The hamming distance 1 between the amino and keto base pairs U,G and A,C are also producing transversions, but here the definite chemical/biological properties remain intact although the interchanges take place between different structures. The base pair U and G follows wobble’s base pairing rule and are critical for the proper translation of genetic rule. Pairing between same bases ((U,U),(A,A), (C,C), (G,G)) have hamming distances 0 as they have exactly same structures and can make proper translation of genetic code. TT and AA play major roles in making water soluble and membrane proteins respectively.

The dual nucleotides occupy the first two positions and thus code for codons along with a nucleotide any one from A/T/C/G at third position, where each codon codes for a specific amino acid. The following diagram (figure 2) gives a schematic representation of the pair of bases to form dual nucleotides in a codon.

**Fig 2:**
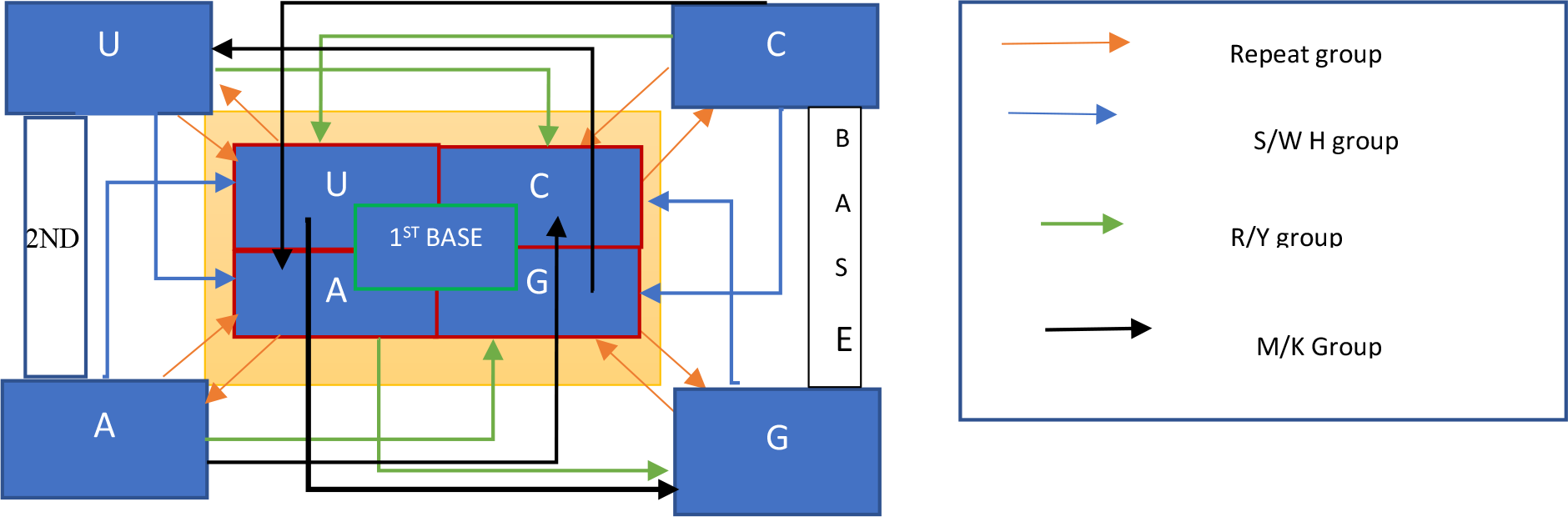
A Schematic Representation of Physicochemical Properties of DNA sequence.

**Table 3:**
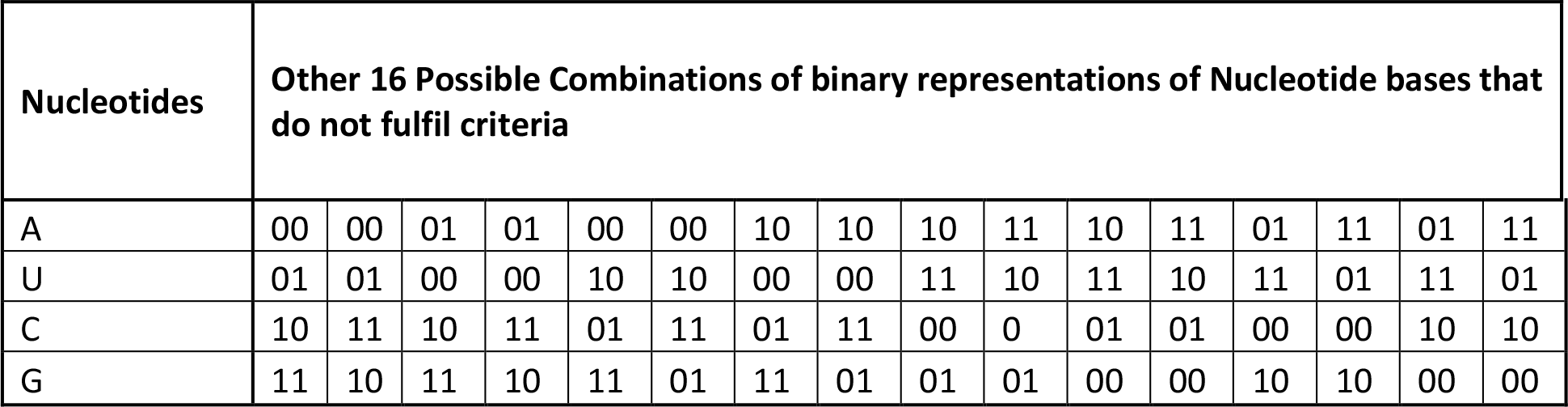
Other 16 Possible Combinations of binary representations of Nucleotide bases that do not fulfil criteria

| Nucleotides | Other 16 Possible Combinations of binary representations of Nucleotide bases that do not fulfil criteria |  |  |  |  |  |  |  |  |  |  |  |  |  |  |  |
| --- | --- | --- | --- | --- | --- | --- | --- | --- | --- | --- | --- | --- | --- | --- | --- | --- |
| A | 00 | 00 | 01 | 01 | 00 | 00 | 10 | 10 | 10 | 11 | 10 | 11 | 01 | 11 | 01 | 11 |
| U | 01 | 01 | 00 | 00 | 10 | 10 | 00 | 00 | 11 | 10 | 11 | 10 | 11 | 01 | 11 | 01 |
| C | 10 | 11 | 10 | 11 | 01 | 11 | 01 | 11 | 00 | 0 | 01 | 01 | 00 | 00 | 10 | 10 |
| G | 11 | 10 | 11 | 10 | 11 | 01 | 11 | 01 | 01 | 01 | 00 | 00 | 10 | 10 | 00 | 00 |

### 1.2 6 bit representation of 64 codons in genetic code

The encoding rule adopted here for the nucleotide bases can encode each codon of genetic code table using 6 bit identifiers accordingly. The first 4 bit of each code carries the binary expressions of physicochemical properties of DNA sequence from which the amino acid sequence is translated and the last two bits are for the third position of a codon which is necessary for small conformational adjustments that affect the overall pairing geometry of anticodons of tRNA.

**Table4:** 6 bit representation of 64 codons in genetic code

|  | U |  |  | C |  |  | A |  |  | G |  |  |
| --- | --- | --- | --- | --- | --- | --- | --- | --- | --- | --- | --- | --- |
| U | UU | U | 000000 | UC | U | 000100 | UA | U | 001100 | UG | U | 001000 |
|  |  | C | 000001 |  | C | 000101 |  | C | 001101 |  | C | 001001 |
|  |  | A | 000011 |  | A | 000111 |  | A | 001111 |  | A | 001011 |
|  |  | G | 000010 |  | G | 000110 |  | G | 001110 |  | G | 001010 |
| C | CU | U | 010000 | CC | U | 010100 | CA | U | 011100 | CG | U | 011000 |
|  |  | C | 010001 |  | C | 010101 |  | C | 011101 |  | C | 011001 |
|  |  | A | 010011 |  | A | 010111 |  | A | 011111 |  | A | 011011 |
|  |  | G | 010010 |  | G | 010110 |  | G | 011110 |  | G | 011010 |
| A | AU | U | 110000 | AC | U | 110100 | AA | U | 111100 | AG | U | 111000 |
|  |  | C | 110001 |  | C | 110101 |  | C | 111101 |  | C | 111001 |
|  |  | A | 110011 |  | A | 110111 |  | A | 111111 |  | A | 111011 |
|  |  | G | 110010 |  | G | 110110 |  | G | 111110 |  | G | 111010 |
| G | GU | U | 100000 | GC | U | 100100 | GA | U | 101100 | GG | U | 101000 |
|  |  | C | 100001 |  | C | 100101 |  | C | 101101 |  | C | 101001 |
|  |  | A | 100011 |  | A | 100111 |  | A | 101111 |  | A | 101011 |
|  |  | G | 100010 |  | G | 100110 |  | G | 101110 |  | G | 101010 |

The chemical features of 20 amino acids have basically depended on eight chemical properties [7],[16],[17]of them. A protein structure is dependent upon the chemical properties of amino acid by which it is formed. So, 20 amino acids are clustered or classified into 8 groups accordingly as shown in following Table 5.

**Table5:** Classification of 20 amino acids according to their chemical properties.

| Group Name | Amino Acids included |
| --- | --- |
| Acidic | Aspartate (D), Glutamate (E) |
| Basic | Arginine (R), Histidine (H), Lysine (K) |
| Aromatic side chain | Tyrosine (Y), Phenylalanine (F), Tryptophan (W) |
| Aliphatic side chain | Isoleucine (I), Leucine (L), Valine (V), Alanine (A), Glycine (G) |
| Cyclic | Proline (P) |
| Sulfur containing | Methionine (M), Cysteine (C) |
| Hydroxyl containing | Serine (S), Threonine (T) |
| Acidic amide | Glutamine (Q), Asparagine (N) |

### 1.3 Mapping between physicochemical properties of DNA sequence and chemical properties of amino acid

When a DNA sequence is translated into primary protein sequence, it finds itself with chemical properties of 20 amino acids. Thus, the nucleotide bases can map their physicochemical properties with the chemical properties of 20 Amino Acids. Mapping between DN Chemical properties and AA Chemical properties can be shown as follows in figure 3:

**Fig 3:**
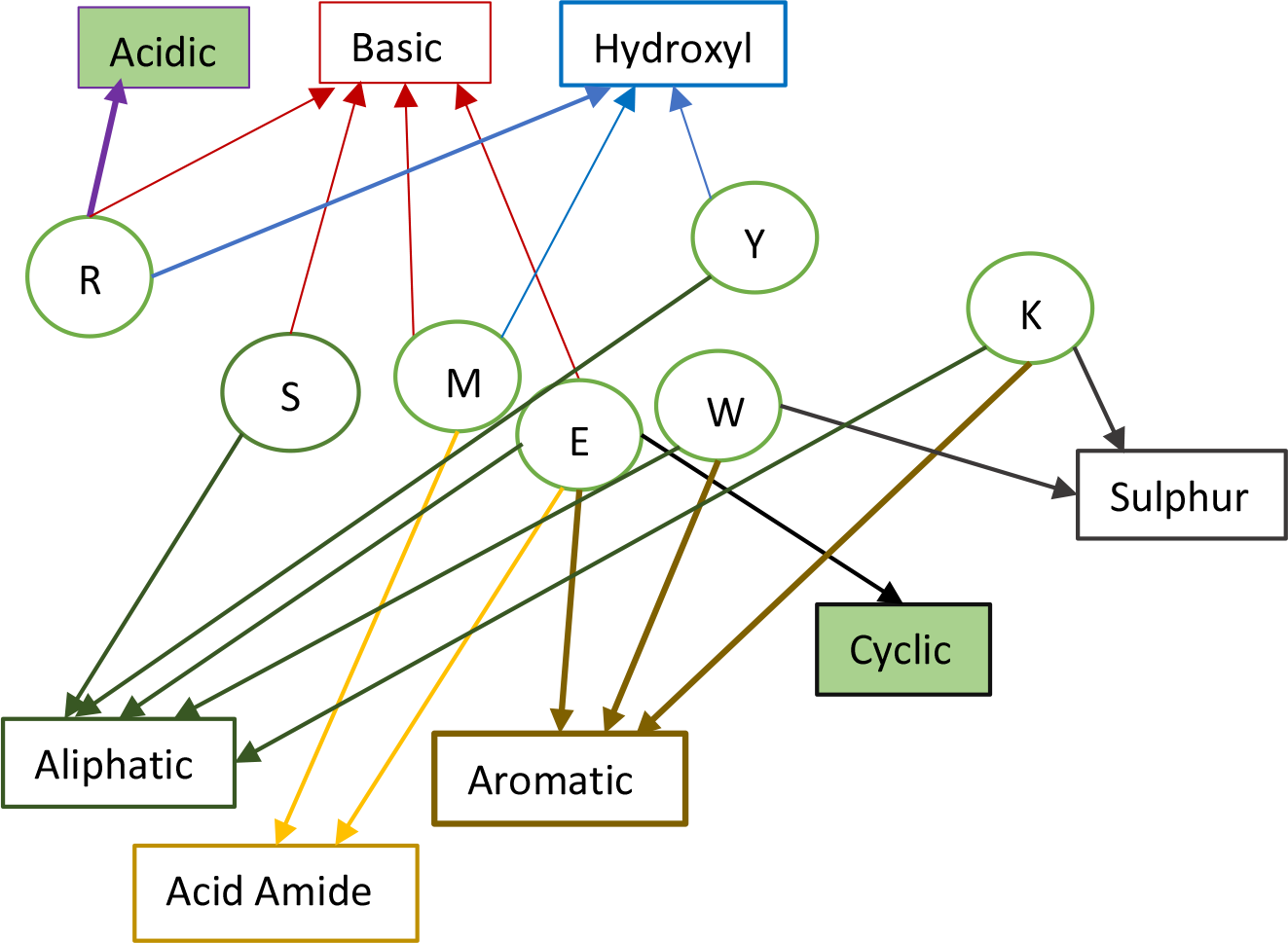
Mapping between physicochemical properties of DNA sequence and chemical properties of amino acid.

The following 7X8 matrix representation specifies the mapping between physicochemical properties of an arbitrary DNA sequence and the chemical properties after translating into Amino acid sequence.

**Table6:** Mapping between physicochemical properties of DNA sequence and chemical properties of its amino acid sequence.

|  | **Acidic** | **Basic** | **Aromatic** | **Aliphatic** | **Cyclic** | **Sulfur** | **Hydroxyl** | **Acid Amide** |
| --- | --- | --- | --- | --- | --- | --- | --- | --- |
| **R** | 1 | 1 | 0 | 0 | 0 | 0 | 1 | 0 |
| **Y** | 0 | 0 | 0 | 1 | 0 | 0 | 1 | 0 |
| **S** | 0 | 1 | 0 | 1 | 0 | 0 | 0 | 0 |
| **W** | 0 | 0 | 1 | 1 | 0 | 1 | 0 | 0 |
| **M** | 1 | 1 | 0 | 0 | 0 | 0 | 1 | 0 |
| **K** | 0 | 0 | 1 | 1 | 0 | 1 | 0 | 0 |
| **E** | 0 | 1 | 1 | 1 | 1 | 0 | 0 | 1 |

Thus, each of 64 codons of genetic table codes for specific amino acid. Each amino acid carries specific chemical property and enriched with physicochemical properties of a DNA sequence from which it is translated. Thus, 20 amino acids can be classified as follows according to chemical properties they contain.

**Table 8:**
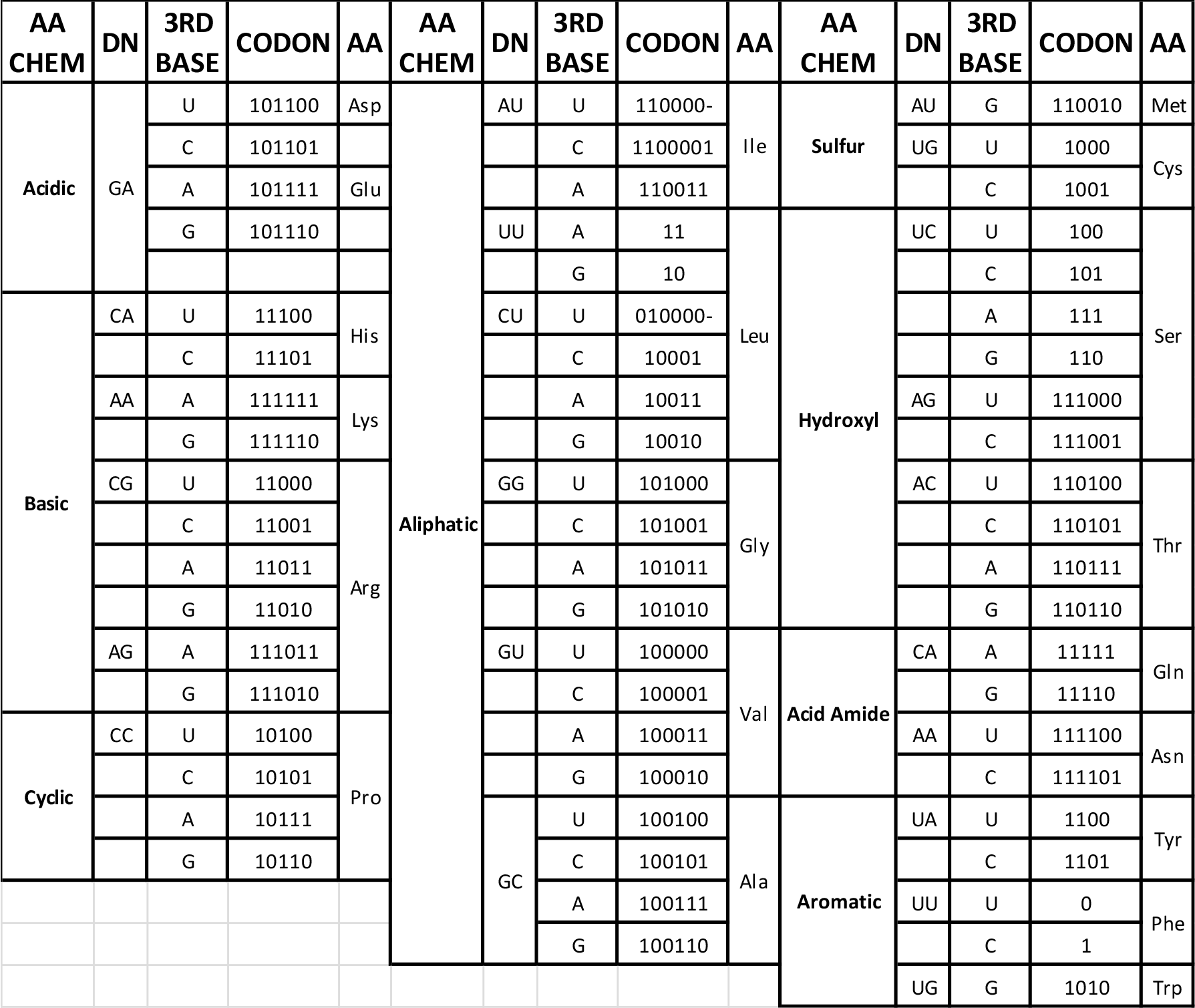
Classification of 20 amino acids according to chemical properties they have and their 6 bit representation.

The classification encodes the mapping between physicochemical properties of DNA sequence and chemical properties of amino acid specified in figure 3 through the light of binary representation.

More microscopic view of the mapping gives us the logic encapsulated between the role of 3 positions of a codon and the final derivation of the amino acid having specific chemical structure. The microscopic view is stated in the figure 4 below. It is to be noted that, in each graph the first layer indicates the physicochemical properties of dual nucleotides and the second layer, the third position of a codon and finally 3^rd^ layer, the chemical properties of 20 amino acids.

**Fig 4:**
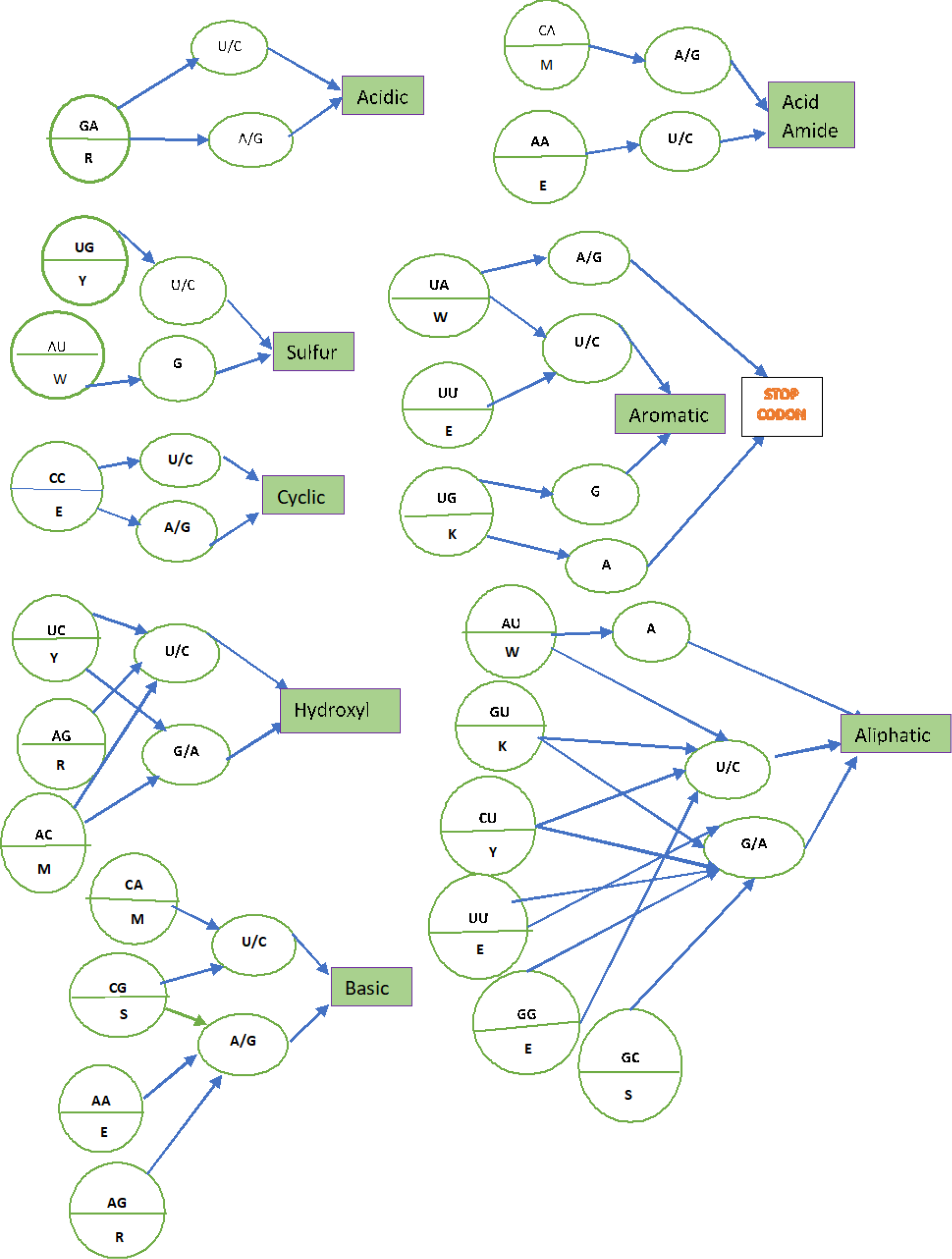
Microscopic view of the mapping between physicochemical properties of DNA sequence and chemical properties of amino acid.

### 1.4 Binary decision tree structure of genetic code table

The Binary decision tree structure of genetic code table mentioned at figure 5 describes each node, which gives the IUPAC abbreviation for the codons consistent with the index chosen. It is worth to mention that, unlike the paper of Nemzer [3], in this manuscript the binary tree has followed the order of the nucleotides of any codon same, i.e.1,2,3. As an example, path leading to methionine, which is coded by AUG is shown in the following table 9.

**Fig 5:**
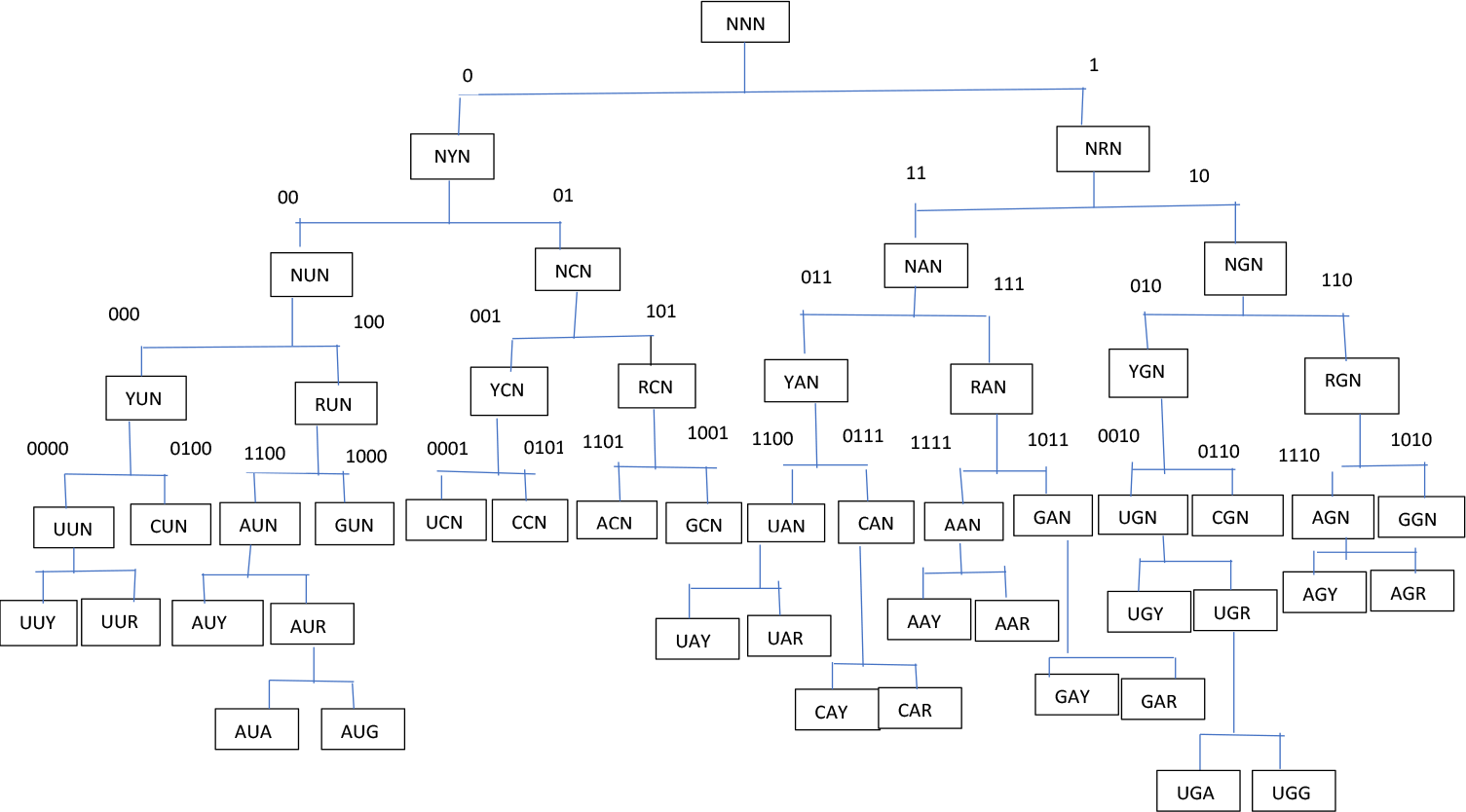
Binary decision tree structure of genetic code table

**Table 9:** Pathway of AUG, in binary tree

| Level | IUPAC | Index |
| --- | --- | --- |
| 1 | NNN | . |
| 2 | NYN | 0 |
| 3 | NUN | 00 |
| 4 | RUN | 100 |
| 5 | AUN | 1100 |
| 6 | AUR | 11001 |
| 7 | AUG | 110010 |

#### Searching codon elements of a DNA sequence from Binary Decision Tree

Now the arbitrary DNA sequence can be read in terms of codon elements using the following algorithm:

1. Start.
2. Make binary representation of the DNA sequence given, where If n=U Then U=00; Else if n=A Then A=11; Else if n=C Then A=01; Else if n=G Then A=10;
3. read a DNA sequence as codons, such that codon sequence C = c1, c2, c3…cj, where j= [n/6] and ci is a codon for any i ∈ {1, 2,…, j}.
4. Start searching procedure,
5. Initialize NNN
6. Go to level 2: If C=NYN Then n=0; Else if C=NRN Then C=1;
7. Go for level 3: If L=NUN; Then C=00; Else if C=NAN Then C=11; Else if C=NCN Then C=01; Else if C=NGN Then C=10;
8. Go to Level4: If L=RUN; Then C=100; Else if C=RAN Then C=111; Else if C=RCN Then C=101; Else if C=RGN Then C=110; If L=YUN; Then C=000; Else if C=RAN Then C=011; Else if C=RCN Then C=001; Else if C=RGN Then C=110;
9. Go to level 5 If L=AUN Then C=1100; Else if L=UUN Then C=0000 Else if L=CUN Then C=0100 elseif L=GUN then C=1000 else if L=AAN Then C=1111; Else if L=UAN Then C=0011 Else if L=CAN Then C=0111 elseif L=GAN then C=1011 else if L=ACN Then C=1101; Else if L=UCN Then C=0001 Else if L=CCN Then C=0101 elseif L=GCN then C=1001 else if L=AGN Then C=1110; Else if L=UGN Then C=0010 Else if L=CGN Then C=0110 elseif L=GAN then C=1010
10. Go to level6 If L=AUY Then C=11000; Else if L=UUY Then C=00000 Else if L=CUY Then C=01000 elseif L=GUY then C=10000 else if L=AAY Then C=11110; Else if L=UAY Then C=00110 Else if L=CAY Then C=01110 elseif L=GAY then C=10110 else if L=ACY Then C=11010; Else if L=UCY Then C=00010 Else if L=CCY Then C=01010 elseif L=GCY then C=10010 else if L=AGY Then C=11100; Else if L=UGY Then C=00100 Else if L=CGY Then C=01100 elseif L=GAY then C=10100 If L=AUR Then C=11001; Else if L=UUR Then C=00001 Else if L=CUR Then C=01001 elseif L=GUR then C=10001 else if L=AAR Then C=11111; Else if L=UAR Then C=00111 Else if L=CAR Then C=01111 elseif L=GAR then C=10111 else if L=ACR Then C=11011; Else if L=UCR Then C=00011; Else if L=CCR Then C=01011; elseif L=GCR then C=10011; else if L=AGR Then C=11101; Else if L=UGR Then C=00101 Else if L=CGR Then C=01100 elseif L=GAR then C=10100
11. Go to Level 7 If L=AUU Then C=110000; If L=AUC Then C=110001; Else if L=UUU Then C=000000 Else if L=UUC Then C=000001 Else if L=CUU Then C=010000 Else if L=CUC Then C=0100001 elseif L=GUU then C=100000 elseif L=GUC then C=100001 else if L=AAU Then C=111100; else if L=AAC Then C=111101; Else if L=UAU Then C=001100 Else if L=UAC Then C=001101 Else if L=CAU Then C=011100 elseif L=GAC then C=101101 else if L=ACU Then C=110100; else if L=ACC Then C=1101001; Else if L=UCU Then C=000100 Else if L=UCC Then C=000101 Else if L=CCU Then C=010100 Else if L=CCC Then C=010101 elseif L=GCU then C=100100 elseif L=GCC then C=100101 else if L=AGU Then C=111000; else if L=AGC Then C=111001; Else if L=UGU Then C=001000 Else if L=UGC Then C=001001 Else if L=CGU Then C=011000 Else if L=CGC Then C=011001 elseif L=GAU then C=101000 elseif L=GAC then C=101001 If L=AUA Then C=110011; If L=AUC Then C=110010; Else if L=UUA Then C=000011 Else if L=UUG Then C=000010 Else if L=CUA Then C=010011 Else if L=CUG Then C=010010 elseif L=GUA then C=100011 elseif L=GUG then C=100010 else if L=AAA Then C=111111; else if L=AAG Then C=111110; Else if L=UAA Then C=001111 Else if L=CAG Then C=011110 elseif L=GAA then C=101111 elseif L=GAG then C=101110 else if L=ACA Then C=110111; else if L=ACG Then C=110110; Else if L=UCA Then C=000111; Else if L=UCG Then C=000110; Else if L=CCA Then C=010111; Else if L=CCG Then C=010110; elseif L=GCA then C=100111; elseif L=GCG then C=100110; else if L=AGA Then C=111011; else if L=AGG Then C=111010; Else if L=UGA Then C=001011 Else if L=UGG Then C=001010 Else if L=CGA Then C=011001 Else if L=CGG Then C=011000 elseif L=GAA then C=101001 elseif L=GAG then C=101000 Exit;

## 2. Application and Results

To validate our methods, PPCA protein family and its four homologs PPCB, PPCC, PPCD, PPCE are taken into account. Periplasmic c7 type cytochrome A (PpcA) protein is determined in Geobacter sulfur reducens along with its other four homologs (PpcB E). From the crystal structure view point the observation emerges that PpcA protein can bind with Deoxycholate (DXCA) [20 21], while its other homologs do not. DNA sequences of PPCA, PPCB, PPCC, PPCD and PPCE are taken from NCBI gene bank.

### 2.1 Find Amino Acid distribution in PPCA and its homologs

Here in this section of the paper the distribution of 20 amino acids in each DNA sequences are investigated. It is worth to state that, PPCA and its 3 homologs PPCB, PPCD and PPCE consist of largest numbers of Lysine (Lys), which has chemical property basic by nature. In other hand PPCC is enriched with Glycine, which is aliphatic by nature.

**Fig 6:**
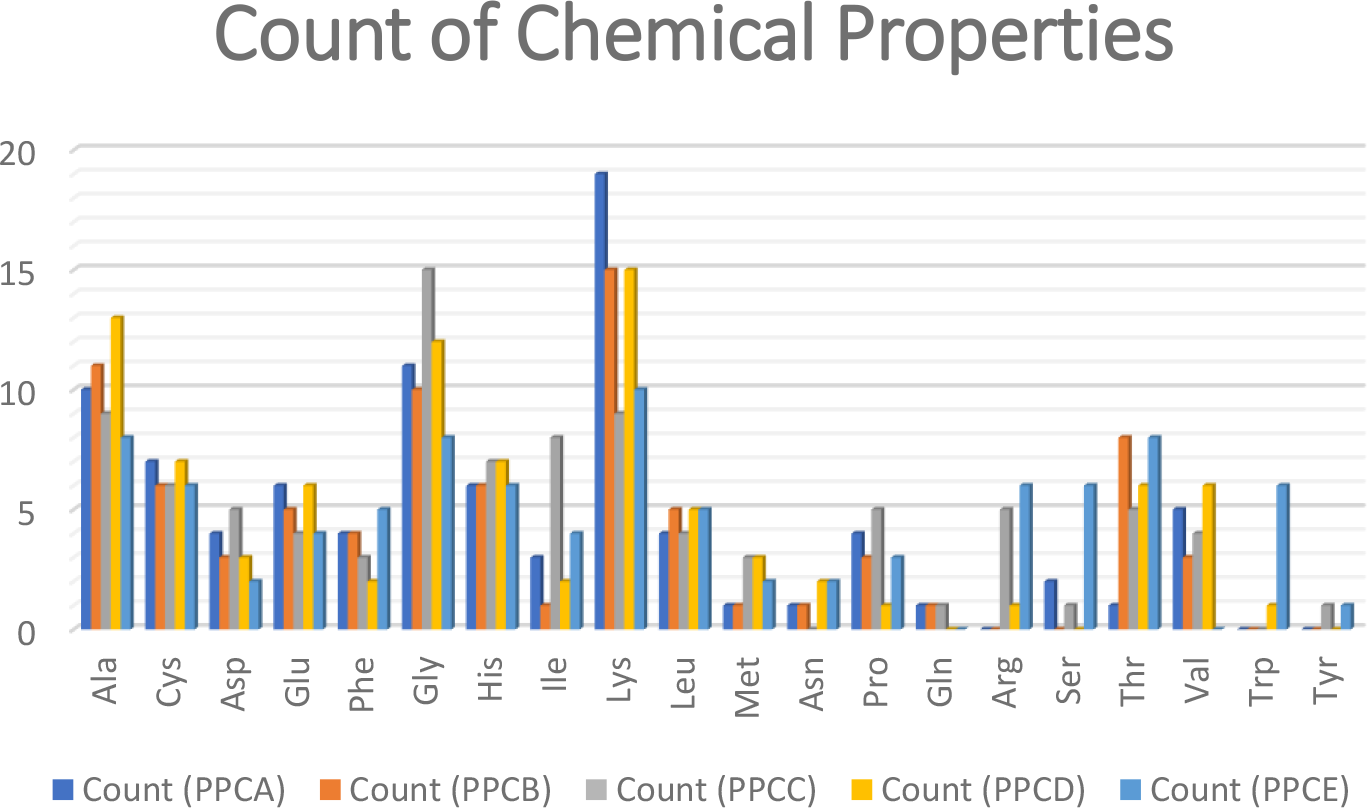
Pictorial analysis of chemical properties present in all the DNA sequences taken.

### 2.2 Make dissimilarity analysis of the DNA Sequences according to chemical group they have

Chemical group distribution test has been carried out for all the DNA sequences taken. It is worth to state that as they are of different lengths, so percentage wise chemical group count has been conducted.

According to figure 7, PPCA and its homologs all have amino acids mostly from aliphatic group. Secondly, they are from basic and thirdly from acidic group. It is remarkable that Aliphatic R groups are hydrophobic and nonpolar, those are one of the major driving forces for protein folding, give stability to globular or binding structures of protein, whereas, Basic and acidic Amino Acids are essential in the formation of Beta sheets, which are secondary protein structures that are stabilized by the creation of hydrogen bonds between acidic and basic amino acids[18]. Physicochemical properties of proteins always guide to determine the quality of the protein structure[18].

**Fig 7:**
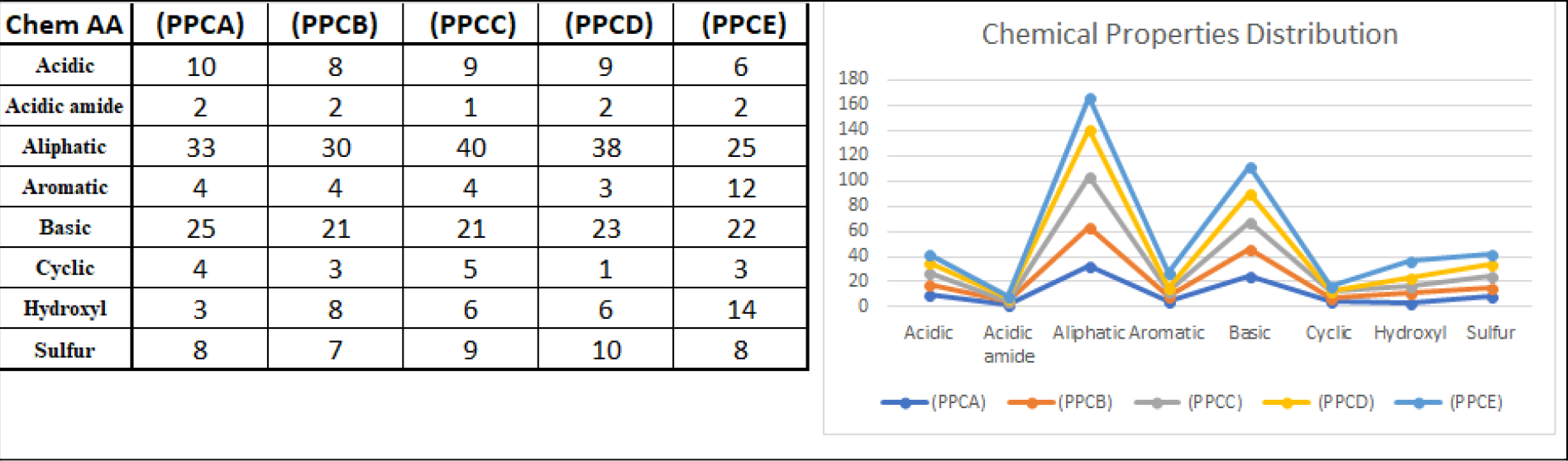
Distribution of chemical groups in each DNA sequences.

Here, 8X1 matrix has been constructed for each DNA sequences as shown as a whole in table 10. Now distance matrix has been constructed to get dissimilarity scores they have among themselves, which is described in table 10 below.

**Table 10:** finding chemical properties percentage wise in all the data set taken.

| Chem AA | PPCA | Chem AA | PPCC | Chem AA | PPCE |
| --- | --- | --- | --- | --- | --- |
| Acidic | 11.23596 | Acidic | 9.473684 | Acidic | 6.521739 |
| Acidic amide | 2.247191 | Acidic amide | 1.052632 | Acidic amide | 2.173913 |
| Aliphatic | 37.07865 | Aliphatic | 42.10526 | Aliphatic | 27.17391 |
| Aromatic | 4.494382 | Aromatic | 4.210526 | Aromatic | 13.04348 |
| Basic | 28.08989 | Basic | 22.10526 | Basic | 23.91304 |
| Cyclic | 4.494382 | Cyclic | 5.263158 | Cyclic | 3.26087 |
| Hydroxyl | 3.370787 | Hydroxyl | 6.315789 | Hydroxyl | 15.21739 |
| Sulfur | 8.988764 | Sulfur | 9.473684 | Sulfur | 8.695652 |

| Chem AA | PPCB | Chem AA | PPCD |
| --- | --- | --- | --- |
| Acidic | 13.5373 | Acidic | 9.782609 |
| Acidic amide | 2.707459 | Acidic amide | 2.173913 |
| Aliphatic | 44.67307 | Aliphatic | 41.30435 |
| Aromatic | 5.414918 | Aromatic | 3.26087 |
| Basic | 33.84324 | Basic | 25 |
| Cyclic | 5.414918 | Cyclic | 1.086957 |
| Hydroxyl | 4.061189 | Hydroxyl | 6.521739 |
| Sulfur | 10.82984 | Sulfur | 10.86957 |

According to table 11, PPCC and PPCD are very similar to PPCA with respect to chemical properties they have. PPCC and PPCD are most similar. In this way PPCB is very much dissimilar to PPCE in the case of physicochemical contents they have.

**Table 11:** Dissimilarity Matrix of PPCA E.

|  | PPCA | PPCB | PPCC | PPCD | PPCE |
| --- | --- | --- | --- | --- | --- |
| PPCA | 0 | 20.4819 | 18.4507 | 18.5149 | 40.7913 |
| PPCB |  | 0 | 24.9912 | 25.4824 | 58.0515 |
| PPCC |  |  | 0 | 11.8535 | 41.3273 |
| PPCD |  |  |  | 0 | 41.3044 |
| PPCE |  |  |  |  | 0 |

### 2.3 Construct well defined mapping between physicochemical properties of dual nucleotide and chemical properties of amino acids through their journey

Here in this part of the experiment, it is tried to show the journey of each DNA sequence starting with any one of the physicochemical properties and lastly converges to which chemical property of an amino acid. The figure clearly shows that except PPCD, PPCA and its remaining homologs are coming from physicochemical property repeating group (AA) and finally getting chemical property basic.

**Fig 8:**
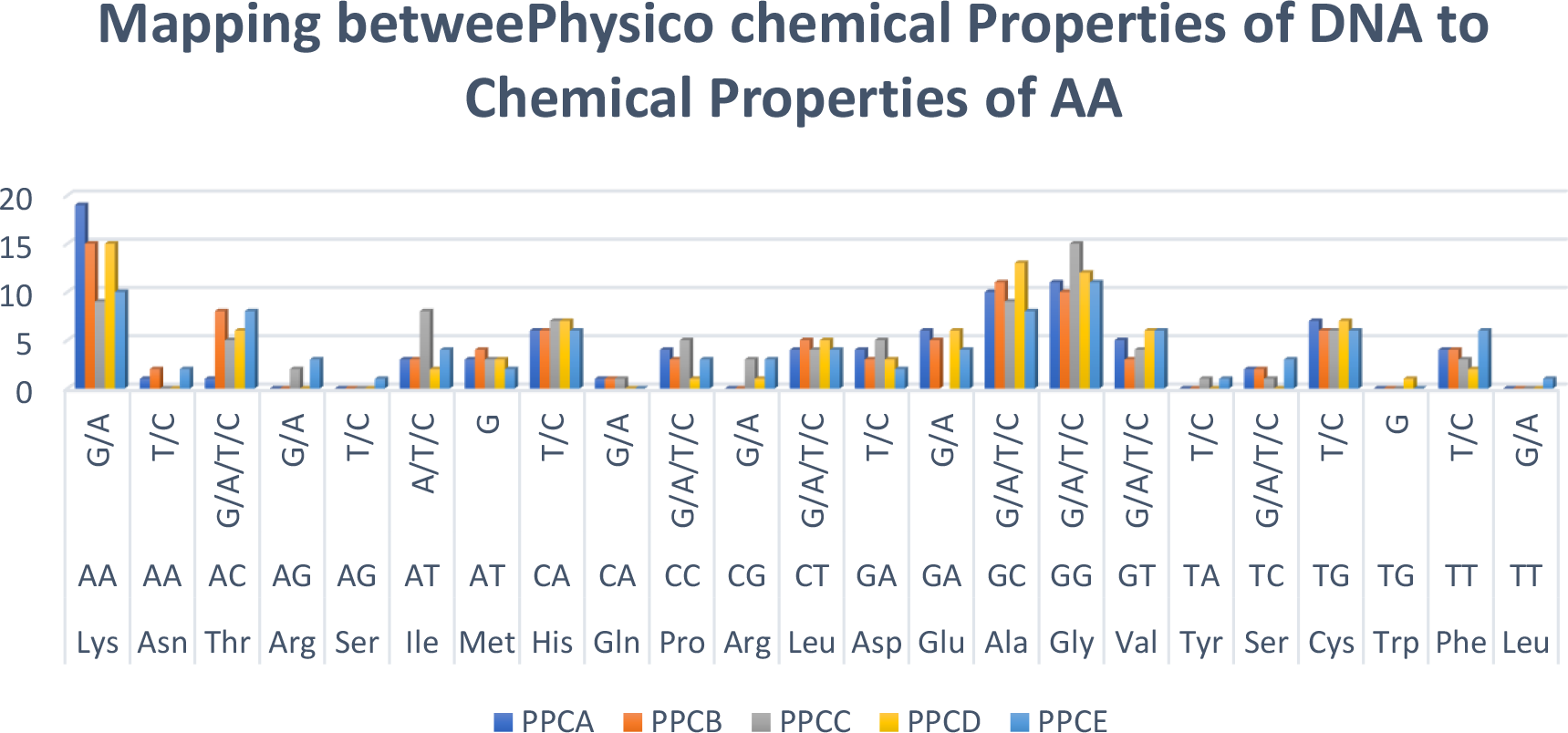
Graphical representation Mapping between Physico chemical Properties of DNA to Chemical Properties of AA.

**Representation of Mapping Between Physicochemical Properties of DNA To Chemical Properties of AA in Terms of Adjacency Matrix.**

### 2.4 Physical properties of protein which are calculated for the data sets

In this section quantitative analysis of some physical properties of amino acid sequences are carried out.

**Table 11:** quantitative analysis of some physical properties of amino acid sequences

| **PPCA E**<br>**Seq No.** | **%**<br>**Access**<br>**Residues** | **%**<br>**Buries**<br>**Residues** | **Hydrophobicity** | **Isoelectric**<br>**Point** | **Molecular**<br>**Weight** |
| --- | --- | --- | --- | --- | --- |
| **PPCA** | 0.010486 | 0.010504 | 0.010486 | 8.31 | 9.75 |
| **PPCB** | 0.010461 | 0.010506 | 0.010493 | 8.23 | 9.67 |
| **PPCC** | 0.010485 | 0.010501 | 0.010484 | 7.96 | 10.08 |
| **PPCD** | 0.010462 | 0.010497 | 0.010488 | 8.1 | 9.7 |
| **PPCE** | 0.010463 | 0.010504 | 0.010488 | 8.86 | 9.9 |

## 3. Conclusion

Transcriptions and translations are the process through which a DNA sequence traverses to get the structure of a primary protein. Many more incidents happen in between, lots of chemistries work. When Crick’s law says about the irreversible flow of information from nucleic acids to proteins, Wobble’s Hypothesis explains reasons for which multiple codons can code for a single amino acid. Irrespective to genes of any families, species, wild type or mutated, our paper here gives a standard model which define a mapping between physicochemical properties of any arbitrary DNA sequence and chemical properties of its amino acid sequence. DNA sequence keeps its signature even after the translation it gets into the properties of an amino acid sequence. The facts and findings are applied on PPCA gene and its four homologs. During experiment it has been observed that when Amino Acid distribution give similar result for PPCA, PPCB, PPCD, PPCE, only PPCC had different observation. It worked the same while mapping between physicochemical properties of dual nucleotide and chemical properties of amino acids has been constructed. Moreover, quantitative analysis of some physical properties of the data set taken finds similarity of PPCA with PPCB, PPCD, PPCE and remarkable difference with PPCC. These findings remark that our standard model of investigation of the journey from DNA to Amino Acid can justify the fact that bio physical properties along with chemical properties of PPCC finds maximum dissimilarities with PPCA. It emerges that, this mapping between physicochemical properties of dual nucleotides and the chemical properties of amino acids provide a new way of looking into the distribution of properties. It is our firm conviction that this investigation will give a new dimension in the task of finding the characteristics of various genes and protein sequences including the diseased and non diseased versions.

